# Genomics of Root System Architecture Adaptation in Sorghum under Nitrogen and Phosphorus Deficiency

**DOI:** 10.1101/2025.05.07.652720

**Authors:** Joshua Magu, Joel Masanga, Catherine Muui, Steven Runo

## Abstract

Plant root systems play a crucial role in water and nutrient uptake so understanding the genetic mechanisms underlying root architecture adaptation to the environment is key, particularly in non-model crops. Here, we investigate the diversity of root system architecture (RSA) in *Sorghum bicolor* under nitrogen (N) and phosphorus (P) deficiency. We used a globally diverse sorghum panel to identify single nucleotide polymorphisms (SNPs) associated RSA responses under N and P deficiency through a genome-wide association study (GWAS). RSA adaptation under P deficiency involved development of fine exploratory roots marked by increased total root length and fine root classes (root length diameter ranges 1 and2), coupled with reduced radial expansion. Under N deficiency, adaptation also involved suppression of root radial growth but without elongation. In both cases, reduced radial growth was marked by reduced surface area and volume, more dramatically for P than N deficiency. GWAS identified SNPs associated with these RSA adaptations, some of which were in regions encoding genes such as: *ILR3-like*, *bHLH*, and a *LEUNIG* homolog all with known roles in root growth regulation. These findings provide novel genetic insights into sorghum root adaptation to nutrient limitations and offer potential targets for breeding resource-efficient crop varieties.

**Summary:** Plants absorb water and nutrients from the soil through their roots, yet for most crops, little is known about how root shape and structure adapt to stressful conditions such as poor soil fertility. In this study, we used a globally diverse collection of sorghum genotypes to investigate how sorghum roots respond to nitrogen and phosphorus deficiency. Specific genotypes showed strong root adaptations under these conditions, leading us to identify genetic factors significantly associated with these responses. Our findings improve our understanding of root adaptation to nutrient stress and highlight promising genetic targets for breeding more nutrient-efficient crops.

## Introduction

Sorghum (*Sorghum bicolor* L. Moench) is a globally important cereal crop used for food, feed, and bioenergy. It can survive adverse conditions such as high temperatures and drought (Trouche et al. 2014), making it a strategic crop for mitigating the impacts of climate change. However, sorghum production in many regions is constrained by poor soil fertility (Khalifa and Eltahir 2023). Specifically, nitrogen (N) and phosphorus (P) are critical determinants of yield (White and Brown 2010); yet their availability is often limited due to depleted soils and the high cost or limited accessibility of fertilizers (Sheahan and Barrett 2017). One way to address nutrient limitations is to exploit plants’ natural ability to adapt their root system architecture (RSA) to optimize nutrient extraction in resource-limited environments.

Previous studies (Parra-Londono et al. 2018) have described three major rooting system types: (i) a small root system, (ii) a compact and bushy root system, and (iii) an exploratory root system, each conferring potential adaptive advantages under nutrient-limited conditions. Genotypes with compact, bushy, and shallow root systems are thought to be better adapted to P scarcity while those with exploratory root systems may be better suited to N scarcity (Parra-Londono et al. 2018). This diversity in RSA underlies strong genotype-dependent responses to nutrient stress. For example, under low P conditions, specific sorghum genotypes respond by increasing root length (Parra-Londono et al. 2018). To offset the metabolic constraints imposed by P deficiency, such plants have reduced secondary growth, resulting in longer, thinner roots (Strock et al. 2017).

Given the importance of these adaptations, this study aimed to identify RSA changes in sorghum under N and P deficiency, as well as the genetic regions that underpin these traits for possible integration into breeding programs targeting nutrient-limited ecosystems.

We hypothesized that sorghum genotypes exhibit extensive genetic variation for RSA traits, based on the species’ ecological adaptations. To test this hypothesis, we used 123 accessions of the sorghum reference set, representing all major cultivated, wild, and intermediate sorghum races from diverse agroecological regions across Africa, India, the Middle East, Europe, and North America (Casa et al. 2008). This panel which comprises mainly landraces, but also contain wild genotypes, breeding material, and advanced cultivars has proven effective for identifying new traits, including parasitic plants (*Striga*) resistance (Kavuluko et al. 2021).

Numerous technologies are now available for root imaging and phenotyping, including hydroponic root elutriation systems (Box et al. 1989), minirhizotron and coring methods (Box et al. 1989; Phillips et al. 2000), non-destructive radar and electrical resistance tomography (Ducut et al. 2022), and root crown phenotyping (Seethepalli et al. 2020). In this study, we used a modified root crown phenotyping approach involving excavation of the upper root system, removal of the growth substrate (vermiculite), and subsequent measurements. The integrated RhizoVision Crown platform (Seethepalli et al. 2020) provided a reliable way to extract data from numerous root systems. Similar platforms have been successfully used in studies of nutrient acquisition in crops such as soybean (*Glycine max*), common bean (*Phaseolus vulgaris*), cowpea (*Vigna unguiculata*), wheat (*Triticum aestivum*), and maize (York 2018).

With advances in high-throughput RNA sequencing (RNA-seq), quantitative trait loci (QTL) can now be resolved at the gene level. In our study, we used genotyping-by-sequencing (GBS) SNP data from the sorghum diversity panel (Morris et al. 2013). This approach has been similarly applied in sorghum (Gelli et al. 2014) to identify genes and novel transcripts involved in RSA control.

To achieve our objective of identifying genetic factors in sorghum that influence root architecture under nutrient-limited conditions, we controlled N and P availability in the growth medium and analyzed the resulting RSA changes, followed by association mapping of these traits with genotype data. We describe these findings and their implications for breeding sorghum varieties better adapted to low-nutrient environments.

## Materials and Methods

### Plant material

A subset comprising 123 accessions of the sorghum reference set [Dataset S1 and (https://genebank.icrisat.org/Common/Viewer?ctg=Referenceset&ref=RS1)] was obtained from the International Crops Research Institute for the Semi-Arid Tropics (ICRISAT) Nairobi, Kenya and maintained at the Plant Transformation Laboratory (Kenyatta University, Nairobi, Kenya).

### Genetic description of sorghum accessions

Sorghum distribution maps were generated using MapTool package (Lemmond, 1994) in R version 4.3.1 based on metadata available at https://www.morrislab.org/data (Lasky et al. 2015). To perform population genetic structure analysis, we retrieved SNP data from the repository of Colorado State University (Morris et al. 2013), available at https://www.morrislab.org/data. Population genetic structure was analyzed using phylogenetic and Bayesian clustering methods.

For phylogenetic analysis, we first converted the retrieved SNP HapMap file into Variant Call Format (VCF) using TASSEL (Trait Analysis by aSSociation, Evolution and Linkage) version 5.0 (Bradbury et al. 2007). The resulting VCF file was then utilized to construct a neighbor-joining (NJ) tree with the ‘ape’ package (Paradis et al. 2004) in R.

For Bayesian clustering, the SNP HapMap file was transformed into PLINK format using TASSEL, then ADMIXTURE software employed to analyze population clusters, evaluating K values ranging from 1 to 10. The optimal number of clusters was determined based on the K value with the lowest cross-validation error. The ADMIXTURE results were visualized using plots generated with the ggplot2 package (Villanueva and Chen 2019) in R.

SNP density was calculated in 50 kb windows using genomic positions extracted from PLINK binary files. Marker quality was assessed based on minor allele frequency and per-marker missing genotype rate. Linkage disequilibrium (LD) was estimated as pairwise *r²* for SNP pairs within the same chromosome and up to 500 kb apart. And LD decay analyzed by binning SNP pairs into 10 kb intervals and computing the mean *r²* per bin for each chromosome and plots were generated using ggplot2 (Villanueva and Chen 2019).

### Sorghum seedling establishment

Sorghum seedlings were sowed in pots measuring 13.5 cm × 11 cm × 11 cm containing vermiculite. For each of the 123 sorghum accessions, 5 seedlings were selected and planted in 3 replicates (5 seedlings per pot in controlled complete randomized) and 3 treatments of watering with; Long Ashton media (Hudson 1967) (optimal N and P), Long Ashton media minus N, and Long Ashton Media minus P. Pots were laid out in a complete randomized block and maintained in the greenhouse for 14 days. Greenhouse conditions were set and maintained at a light/ dark cycle of 12/12 h and a temperature of 28℃ day and 24℃ nights.

### Root imaging and analysis of sorghum phenotypes

On the 14th day, fresh root samples were photographed using a Nikon DX VR 18–140 mm camera at a resolution of 1400 dpi. The acquired images were processed and analyzed using RhizoVision Explorer, (https://www.rhizovision.com/) an open-source software designed for root phenotyping.

Images were firstly scaled and cropped to produce grayscale versions, followed by the application of a global image threshold to distinguish roots (foreground) from the background using the software’s ‘Broken Roots’ mode. The pixel-to-centimeter conversion was calibrated to 41.02 pixels per mm. Image thresholding was set to 150 to achieve clear root boundaries while minimizing external noise. Non-root artifacts smaller than 2 mm² were filtered out to prevent interference with the analysis. Edge smoothing was enabled and set to level 2 to reduce false root appearances during skeletonization, thereby enhancing the delineation of root features. The root pruning threshold was configured to remove false root segments where tips are shorter than the diameter of the primary root, ensuring accurate feature extraction. RhizoVision Explorer output 13 RSA traits, were exported as .csv files (Dataset S2).

### Data analysis

RhizoVision data (Dataset S2) were used for downstream analysis in R version 4.3.1. Hierarchical clustering was performed using the agnes() function from the cluster package (Maechler et al. 2026) using Ward’s method and clusters visualized using dendrograms. PCA was performed on all RSA traits using the FactoMineR package (Lê et al. 2008) and biplots generated with the factoextra package (Kassambara and Mundt 2020). Correlation analyses were performed using the corrplot package (Wei and Simko 2021), and additional statistical analysis with the PerformanceAnalytics package (Peterson 2020). Gaussian graphical models (Glasso networks) were generated using the qgraph package (Epskamp et al. 2012). Node centrality measures were calculated from the Glasso networks using the centrality() function. Linear mixed-effects models were performed using the lme4 package (Bates et al. 2015) and Forest plots generated using ggplot2.

Analysis of variance was performed to evaluate the effect of genotype; effect of environment; and effect of both genotype and environment on root traits. Mean separation was performed using Tukey’s Honest Significant Difference (HSD) test at p ≤ 0.05. Heritability of each trait under the tree conditions was estimated according to (Hill et al. 2008)

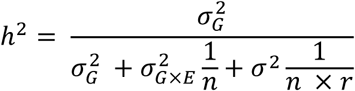

Where 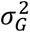 represent the genotypic variance, 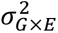 denotes variance of genotype by environment interaction, *σ*^2^ error of variance, n number of environments and r number of replicates.

### Genome wide association studies (GWAS)

Genome-wide association studies (GWAS) were conducted on RSA phenotypes under three conditions: optimal, nitrogen-deficient, and phosphorus-deficient. A total of 264,962 previously developed and filtered single nucleotide polymorphisms (SNPs) (Morris et al. 2013) were used for genome-wide association analysis. GWAS was first conducted on individual RSA traits under each treatment condition, followed by analysis of traits that showed significant associations in a multivariate framework. All analyses were performed using the Genome Association and Prediction Integrated Tool (GAPIT) version 2 (Tang et al. 2016), implementing three models: Bayesian-information and Linkage-disequilibrium Iteratively Nested Keyway (BLINK) (Huang et al. 2018), the Fixed and Random Model Circulating Probability Unification (FarmCPU) (Kusmec and Schnable 2018) and the Multi-Locus Mixed Model (MLMM) (Segura et al. 2012). To correct for multiple testing, we applied a 5% false discovery rate (FDR). The results were visualized using quantile-quantile (QQ) plots and Manhattan plots. Significant SNPs associated with the phenotypic traits were mapped onto the *Sorghum bicolor* genome using Phytozome v3.1.1’s JBrowse tool (Goodstein et al. 2012). Candidate genes of interest were selected based on their proximity to the SNPs and their potential involvement in the regulation of root development.

## Results

### Description of sorghum root system architecture

To extract root data from various sorghum genotypes, we used the RhizoVision platform, which provided 13 root traits. These traits are illustrated in Figure 1 and described in more detail in Table S1. The traits can be grouped as follows: (i) Size: Total root length (TRL), Perimeter (P), Surface Area (SA), Network Area (NA), and Volume (V); (ii) Extent: Branch Points (BP), Branching Frequency (BF), and Number of Root Tips (NRT); (iii) Distribution: Average diameter (AD), Median diameter (MD), and Maximum diameter (MD).

**Figure 1.**
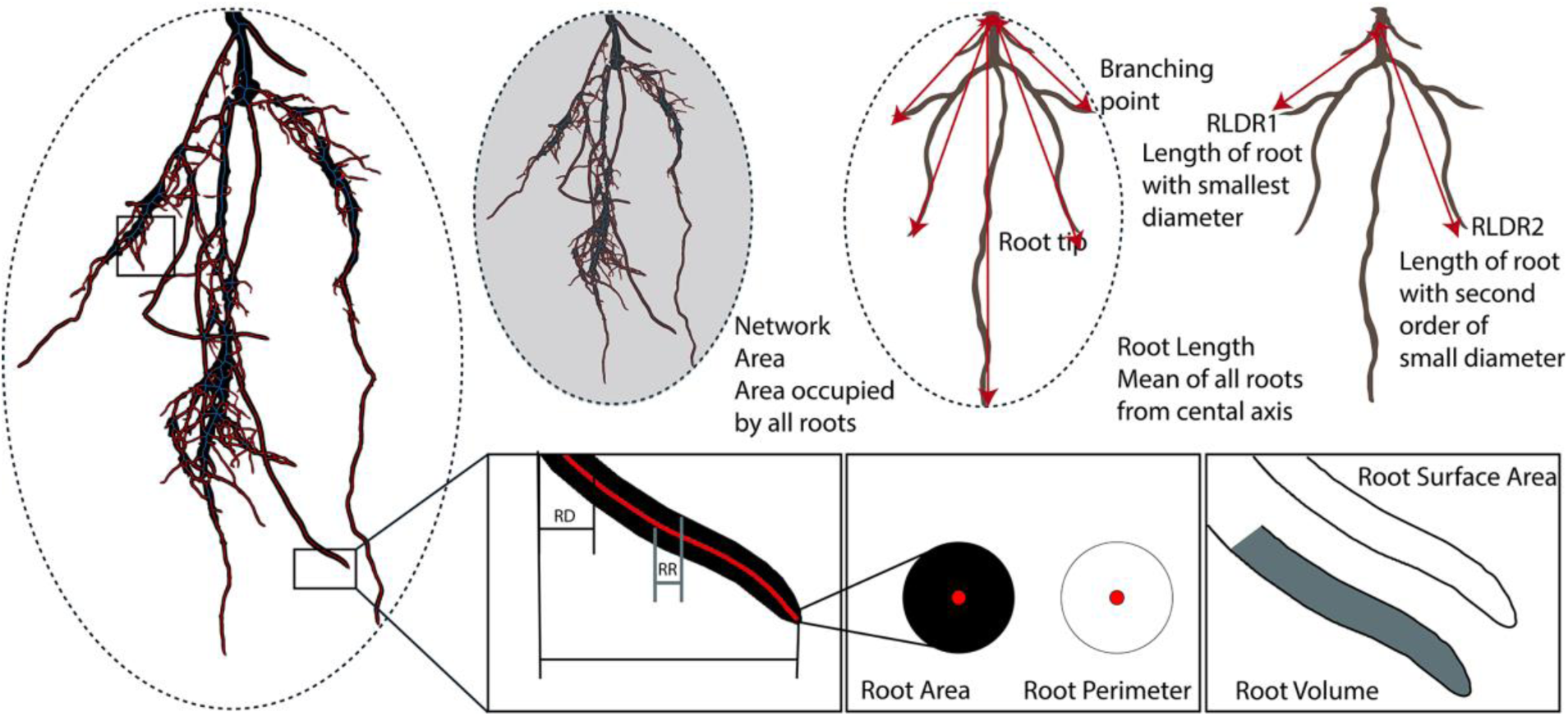
RhizoVision analysis of root system architecture showing a skeletonized sorghum root image used to extract measurements. The upper panel shows measurement for network area (left panel), root length, root tip and branching point (middle panel) and root length diameter range 1 (RLDR1) and RLDR2 (right panel). The lower panel shows a small root region that is amplified to show root diameter (RD), and root radius (RR) in the left panel, root area and root perimeter (middle panel) and root volume and root surface area (right panel). Scale of skeletonized root crown = 140 mm.

### Genetic variation of sorghum root system architecture and response to N and P nutrients deficiency

To examine genetic variation in the sorghum root system we used a subset of 123 genotypes of the sorghum diversity panel (Figure S1a). The panel was highly diverse and comprised all major sorghum races. Phylogenetic analysis using Neighbor-Joining grouped the genotypes into major clades corresponding to races: bicolor, caudatum, durra, guinea, Guinea subraces, kafir and wild genotypes with some clusters showing admixture (Figure S1b). These same groupings were reflected in Bayesian hierarchical clustering (BHC) (Figure S1c) at the most optimal K (K=4 to K=6). Plots from BHC also showed weak population structuring rather than strictly discrete genetic groups. Additionally, the sorghum genotypes represented populations originating in diverse agro-ecological regions across Africa, Asia, America and Europe (Billot et al. 2013).

We then grouped the 123 genotypes used in the study based on Euclidean distances of all their RSA traits. Under optimum nutrient conditions, this grouping produced 5 clusters (Figure 2) but under N and P deficiency, the clusters reduced to 2 suggesting that nutrient deficiency led to changes on the RSA of genotypes leading to reorganized the relationships.

**Figure 2.**
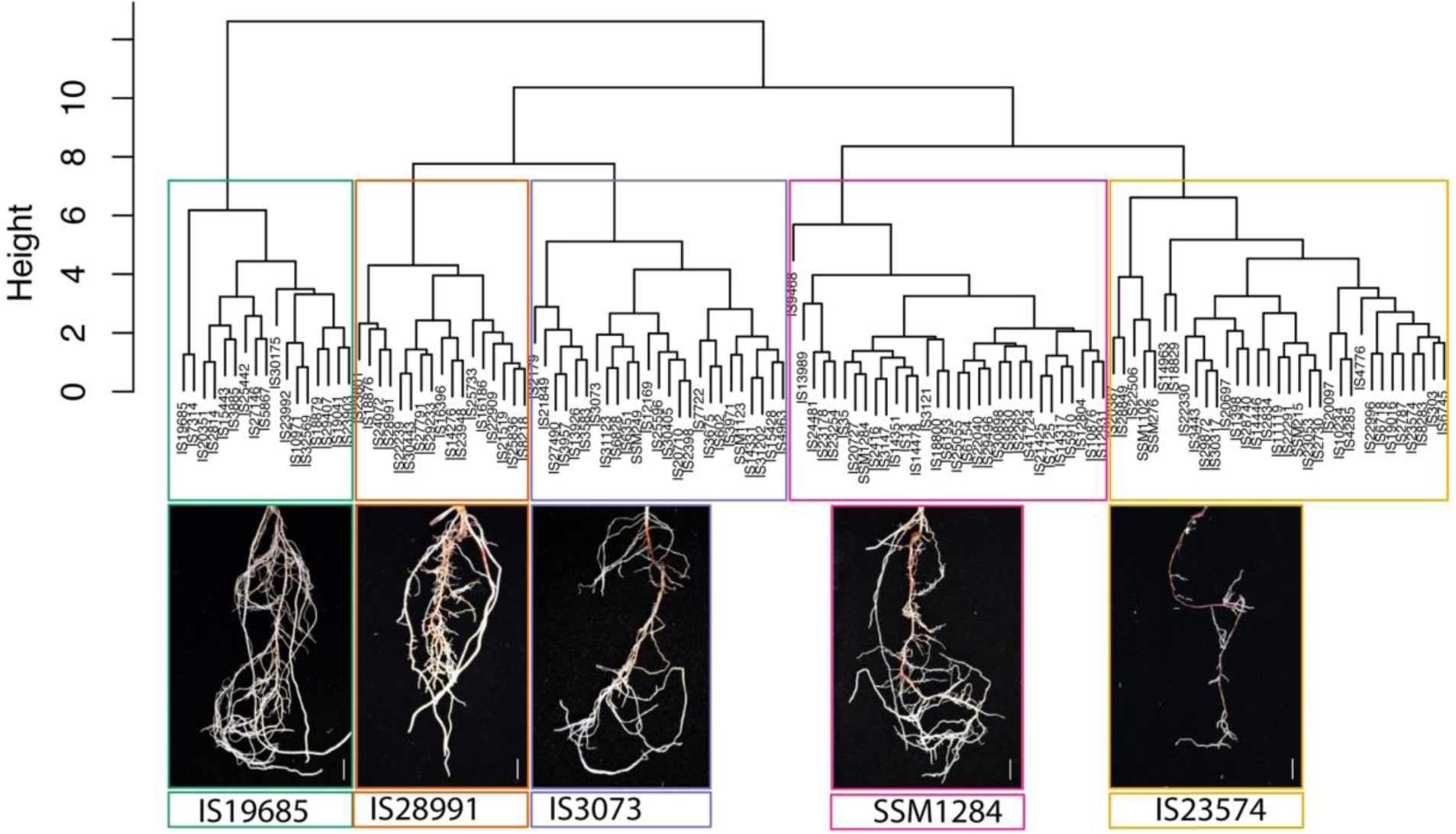
Variation of sorghum root architecture: Dendrogram showing clustering of root traits from a dataset of 123 sorghum genotypes. Clustering was performed using the complete method and Euclidean distance using all analyzed root system architecture traits. The colored boxes highlight five distinct root-type clusters, each represented with crown images corresponding to the genotypes within the cluster.

### Partitioning of genotypic variation of sorghum root system architecture

Next, we examined how variation in RSA was partitioned among genotypes using principal component analysis (PCA) under optimal, minus N, and minus P conditions.

Under optimal nutrient conditions (Figure 3A), the first principal component (PC1; 56.9% of the variance) was mainly associated with total root length, number of root tips, number of branching points, perimeter, surface area, and root length diameter ranges 1 and 2. These traits exhibited long vectors with high contribution and loading values, indicating a strong influence on PC1 (Dataset S4). The second principal component (PC2; 31.5%) was contributed by root diameter traits: maximum diameter, average diameter, and median diameter, which were oriented oppositely to branching frequency implying negative correlation between root thickness and branching frequency.

**Figure 3.**
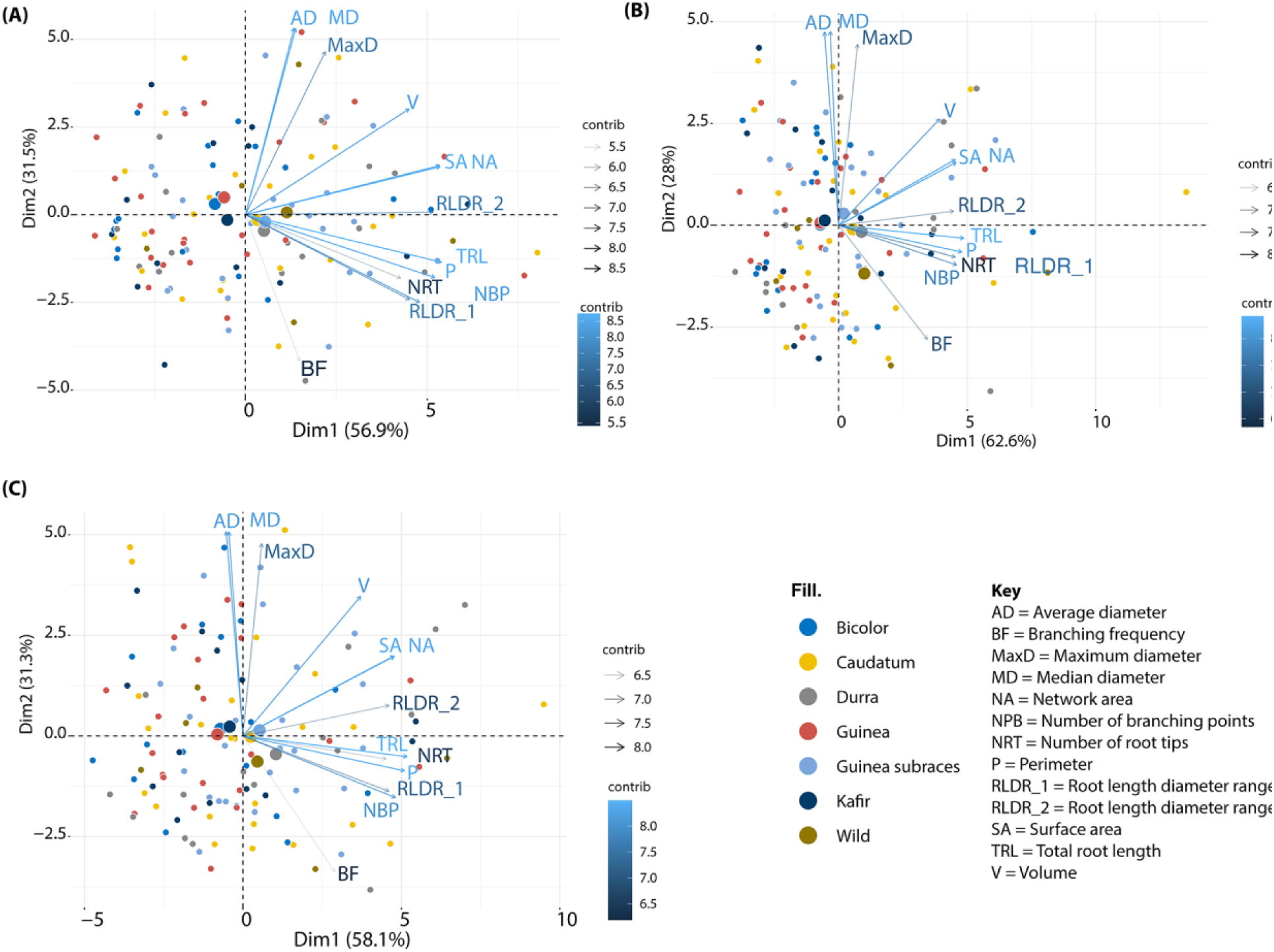
Principal component analysis (PCA) of root system architecture (RSA) traits across nutrient conditions. (A) optimal nutrient conditions, (B) nitrogen deficiency, and (C) phosphorus deficiency. Arrows indicate RSA trait loadings, with arrow length and color intensity reflecting the relative contribution of each trait to the principal components. Dots represent individual genotypes positioned according to their RSA trait profiles. Genotypes projecting in the direction of a given trait arrow exhibit higher values for that trait, whereas genotypes positioned opposite the arrow show lower values. The percentage of variance explained by each principal component is indicated on the axes.

Under N deficiency (Figure 3B), PC1 accounted for 62.6% of the variance contributed by total root length, number of root tips, number of branch points, perimeter and root length diameter ranges 1 and 2. PC2 accounting for 28.0% variance was still associated with diameter-related traits, and branching frequency was also negatively aligned with PC2 like in optimal conditions. RSA vectors were also notably tighter implying that they work as a module.

Under P deficiency (Figure 3C), PC1 explained 58.1% of the variance and remained associated with total root length, number of root tips, number of branch points, perimeter and root length diameter ranges 1 and 2; however, vector alignment among these traits was weaker than under N deficiency. Linke in N deficiency, branching frequency contributed negatively along PC2 under P deficiency.

Together, these patterns indicate that nutrient deficiency enhances adaptive changes in RSA, with diameter (thickness) traits becoming increasingly uncorrelated from elongation and branching traits under both N and P deficient conditions.

Analysis of individual genotypes (dots) on the PCA plots revealed that under optimal conditions, genotypes were widely dispersed, reflecting high internal diversity with respect to all RSAs (Figure 3A). Under N and P deficiency (Figure 3B and C) some genotypes spread primarily along PC1, projecting strongly towards elongation/branching traits, and fewer towards PC2 (diameter and branching frequency RSA) plausibly reflecting adaptability of these genotypes based on the RSAs that they aligned with. However a majority of genotypes were at the center of the PCA, suggesting that RSA traits of these genotypes do not adapt to N or P deficiency.

### Interactions of sorghum root system architecture traits

To determine levels of correlations among various RSA traits under all experimental conditions, we performed a Person’s correlation analysis.

Under optimal conditions, (Figure 4A), analysis revealed that total root length was the central trait showing a strong positive correlation with all RSA traits (r > 0.77) except diameter based traits (non-significant), branching frequency (weak correlation r = 0.38) and root volume (moderate correlation, r = 0.63). In contrast, diameter-related traits; maximum diameter, average diameter, and median diameter were negatively correlated with branching frequency.

**Figure 4.**
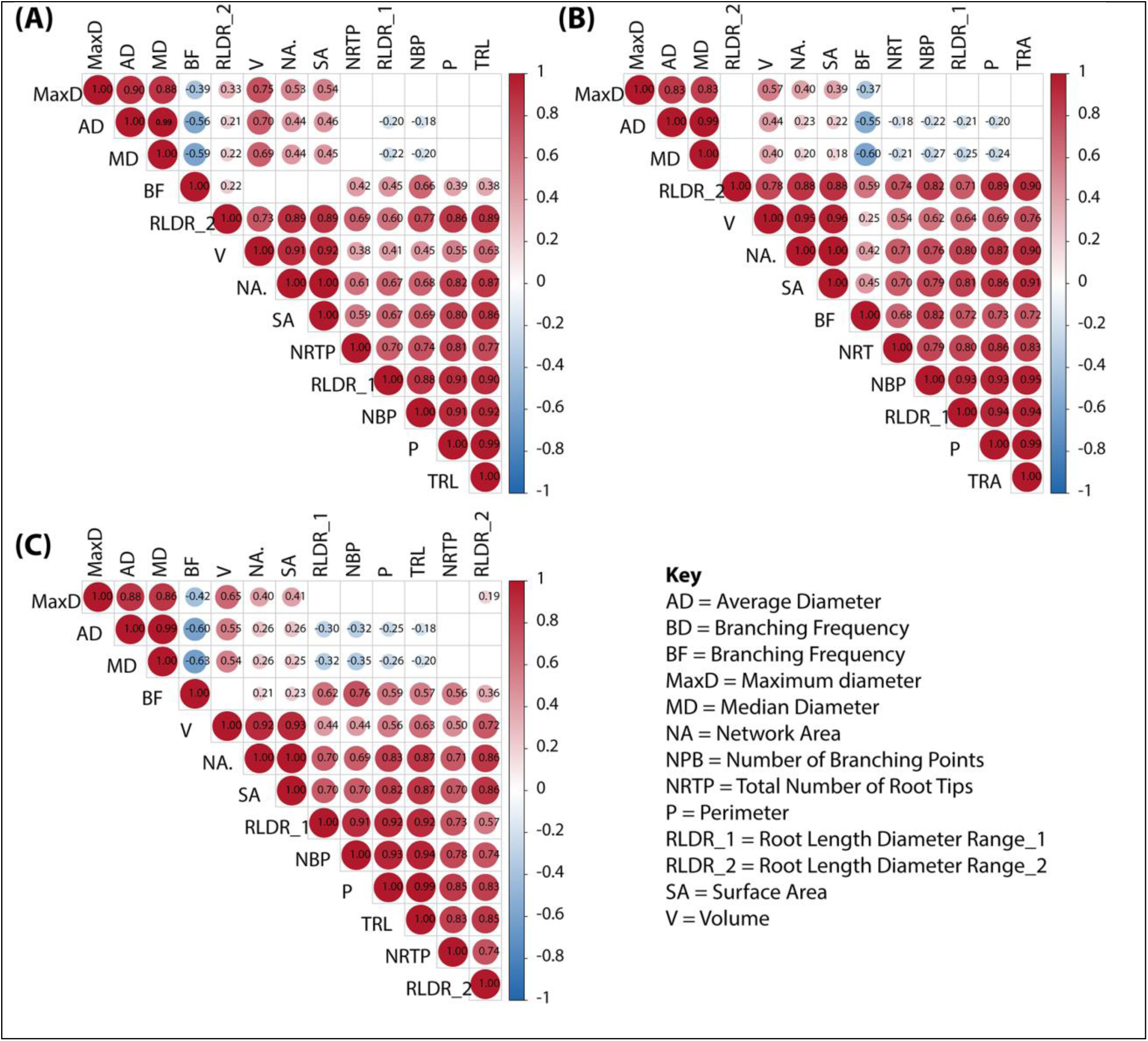
Pearson’s correlation analysis of root system architecture traits under (A) optimal, (B) nitrogen deficiency, and (C) phosphorus deficiency conditions. Colored cells show Pearson correlation coefficients (*r*), with values displayed. Only significant correlations (*P* < 0.05) are shown, and traits are ordered by hierarchical clustering. Traits measured are shown on the key.

Under N deficiency (Figure 4B), total root length showed strong positive correlations (r > 0.7) with all measured RSA traits, except for root diameter traits (maximum, average, and median diameter). Notably, these diameter-related traits exhibited negative correlations with branching frequency.

Under minus P conditions (Figure 4C), root length diameter range 2 was the central trait, showing strong positive associations with most RSA traits (r > 0.7), but no significant correlations with diameter-based traits. Its associations with branching frequency (r = 0.36) and with root length–diameter range 1 (r = 0.57) were weak. Diameter-based traits under low P conditions were more strongly and negatively correlated with branching frequency than under optimal or N-deficient conditions; for example, median diameter showed a stronger negative correlation under low P (r = −0.63) compared with optimal (r = −0.59) and low N conditions (r = −0.60).

These findings indicate that the negative association between root thickness (diameter-based RSA) and branching was consistent across treatments but strongest under P deficiency, highlighting nutrient-specific adaptations of root system architecture and plausible trade-offs between branching and thickness.

Gaussian graphical models (Bhushan et al. 2019) were then used to quantify conditional dependencies among root traits (Figure 5 A-C). Under optimal conditions (Figure 5A), the network was highly integrated, with strong positive partial correlations among total root length, perimeter, network area, surface area, and volume. Diameter traits formed a distinct module that remained connected to overall root size, while branching traits were positively associated with total root length.

**Figure 5.**
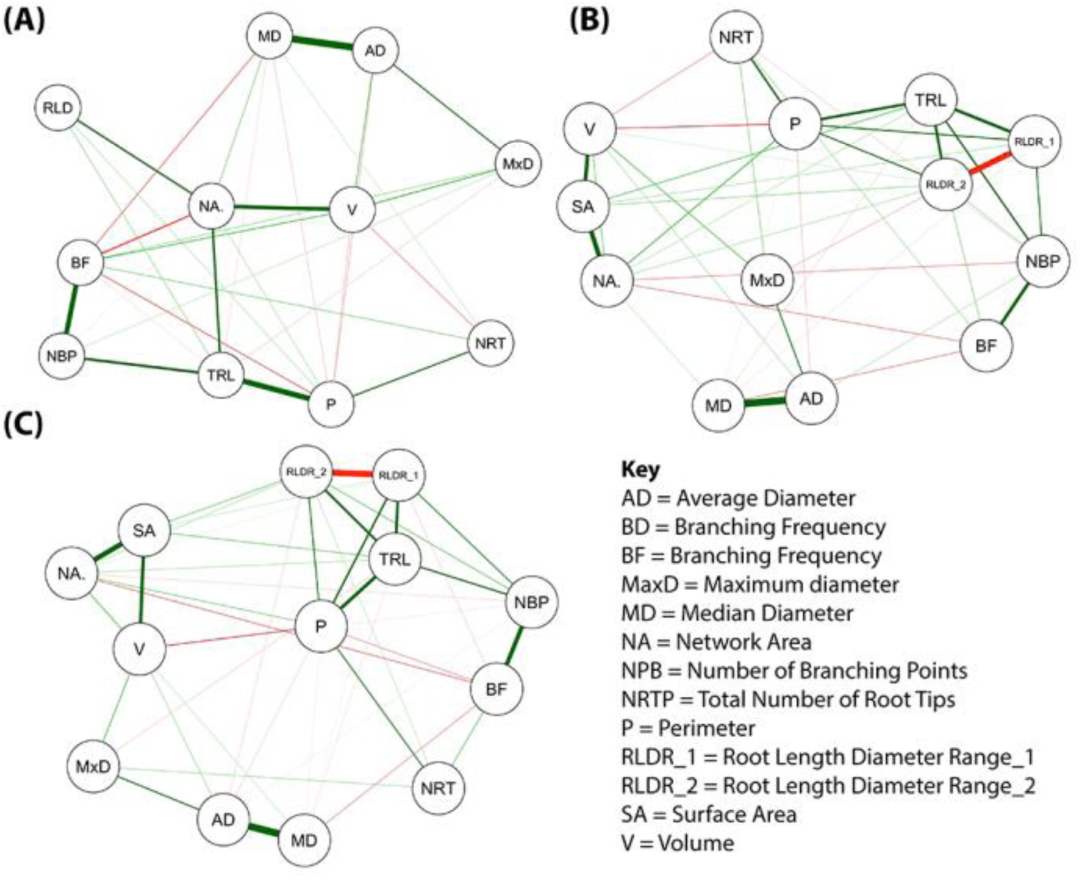
Gaussian graphical model showing strength of correlation between traits: A. Optimal nutrients, B, nitrogen deficiency and C. Phosphorus deficiency. Red and green lines show negative and positive correlations, respectively.

Under N deficiency (Figure 5B), traits clustered into modules, with total root length emerging as a central hub linking branching traits (number of branch points and branching frequency) and root length–diameter range ranges 1 and 2. There was also notable negative correlations involving diameter traits.

Under P deficiency, the network exhibited reduced connectivity, characterized by strong connectivity between total root length and root length diameter ranges 1 and 2. There were also notable negative correlations involving size and diameter traits.

These patterns indicate a shift from coordinated general root growth under optimal conditions to increasingly specialized root system architectural strategies: branched roots under N deficiency, and elongated fine roots under P deficiency with diameter traits acting negatively to the traits.

### Impact of nutrient stress on sorghum root architecture

To assess the response RSA traits to N and P deficiency, we performed fixed-effects model analyses for all traits, using the optimal nutrient treatment as the reference. The results showed that root length–related traits: root length in diameter ranges 1 and 2 and total root length increased markedly under P deficiency. In contrast, RSA traits associated with secondary growth, surface area and volume, were reduced under both N and P deficiency, with a more notable reduction observed under P deficiency (Figure 6; Dataset S3).

**Figure 6.**
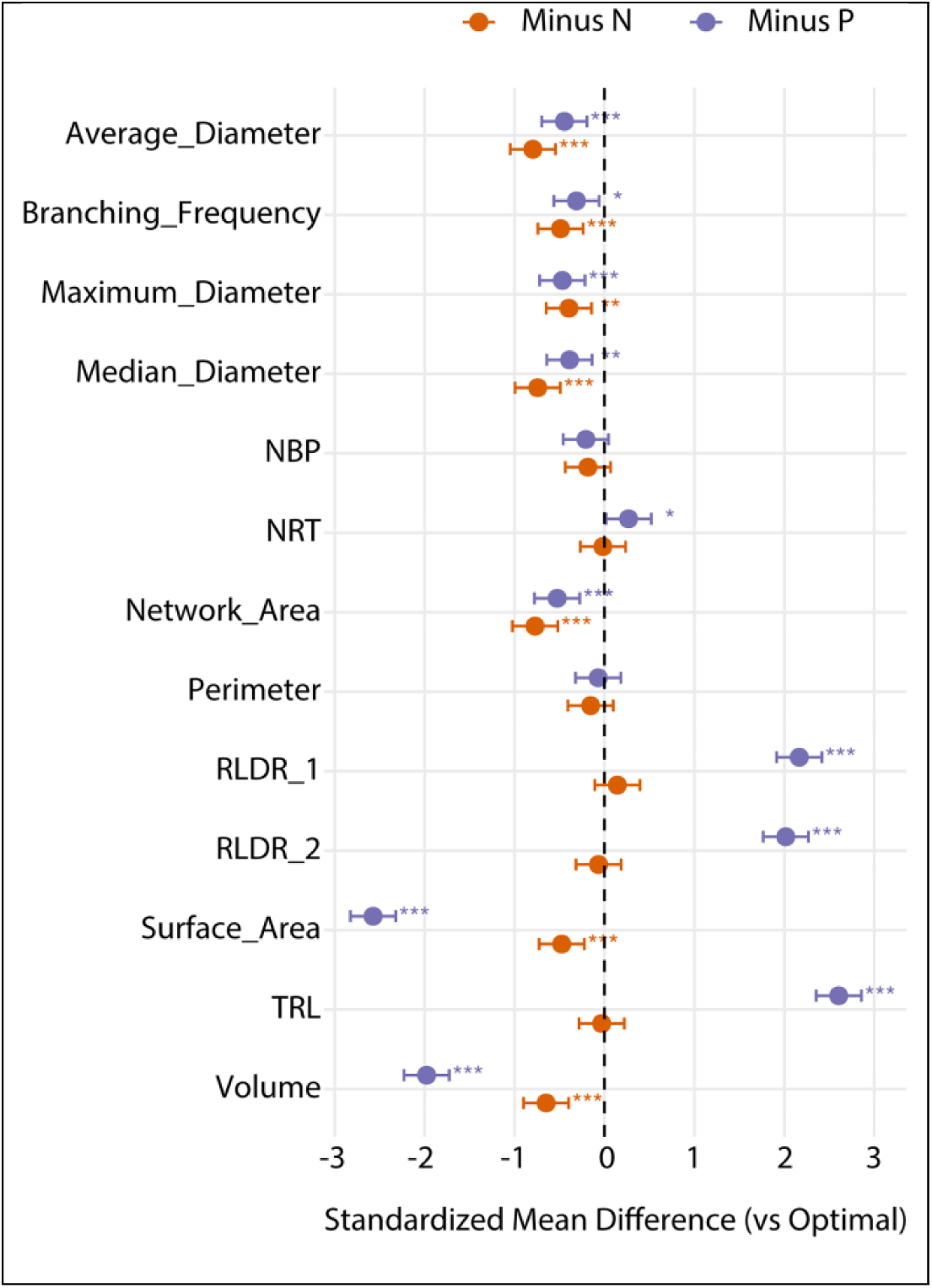
Impact of nutrient stress on sorghum root system architecture. Forest plot showing standardized mean differences (SMD ± 95% confidence intervals) of root system architecture traits under nitrogen and phosphorus deficiency relative to the optimal nutrient treatment (reference = 0). Positive SMD values indicate an increase relative to the optimal treatment, whereas negative values indicate a decrease.

**Figure 7:**
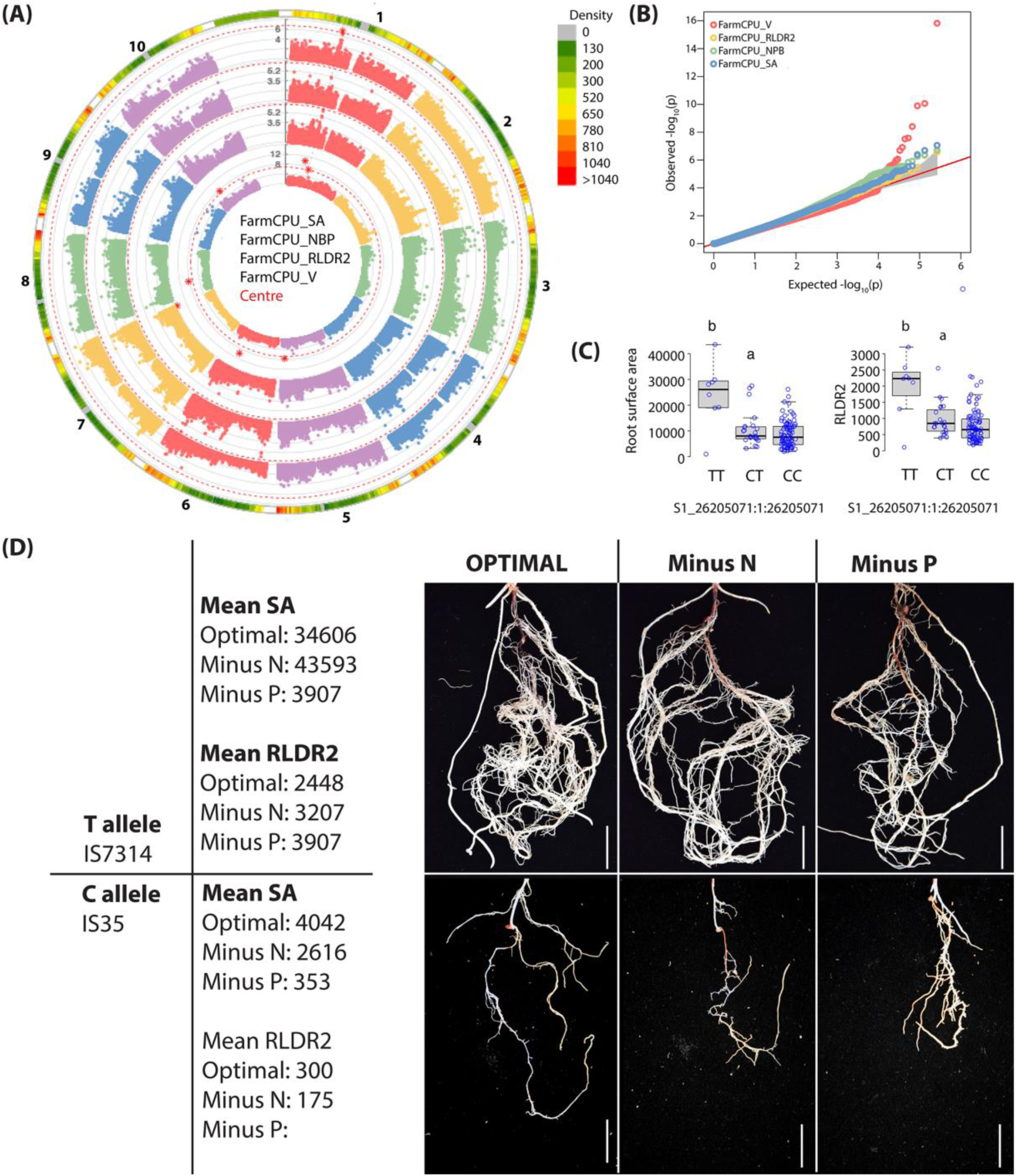
Genome-wide association studies identifying genetic factors influencing root system architecture traits in sorghum. (A) Fixed and Random Model Circulating Probability Unification model circular Manhattan plot highlighting significant single nucleotide polymorphisms (SNP) associated with root system architecture traits, including volume, root length diameter 2, number of branch points, and surface area. (B) Corresponding Quantile-quantile (QQ) plot comparing observed and expected p-values to assess model accuracy and false positives. (C) Boxplot illustrating the allelic effect of the most significant SNP (S1_26205071) on surface area and root length diameter range 2. Favorable alleles increase trait expression, while unfavorable alleles decrease it. (D) Representative root phenotypes comparing “C” and “T” allele genotypes under optimal, nitrogen-deficient, and phosphorus-deficient conditions. Genotypes with the “C” allele exhibited significantly higher SA and RLD2. Scale bar = 140 mm.

Together, these findings indicate that P deficiency promotes an increase in total root length, largely driven by the proliferation of fine roots. This response occurs alongside a reduction in overall root surface area and volume traits under both nutrient limitations, particularly under P deficiency. In all cases, radial growth is suppressed.

### Genome wide association studies to identify genetic factors underpinning root system architecture traits in sorghum

To investigate the genetic factors contributing to the observed RSA responses under N and P stress, we conducted GWAS on all 13 measured traits using GBS based SNP markers. Assessment of markers for this dataset SNP markers showed that they were evenly distributed across the genome with no large gaps (Figure S4). Marker quality metrics indicated low missing rates and balanced minor allele frequencies across chromosomes (Figure S4). And Linkage disequilibrium decayed within the first 100 kb and approached background levels by ∼300–500 kb, (Figure S4).

In total, GWAS detected 25 significant SNPs detected under N and P deficiency using RSA traits of volume, surface area, root length diameter range 2, and number of branching points (N deficiency) and root length diameter range 2 (P deficiency).

The most prominent SNP was S1_26205071 identified 11 times using volume, surface area, number of branching points, and root length diameter range 2, under N deficiency and root length diameter range 2, under P deficiency. This SNP did not fall on a gene region, but its highly significant P value and association with multiple traits in two models pointed to its importance in shaping the root architecture. Therefore, we characterized the phenotypic variation related to each haplotype of the SNP change. We determined that 92 and 8 accessions harbored the C and T haplotypes, respectively, whereas the remaining 37 accessions lacked genotype information. Under all environments, the accessions carrying the T allele showed significantly higher values of the RSA traits surface area, root length diameter range 2, and number of branching points than those carrying the C allele (p ≤ 0.05), indicating that the T haplotype had a positive effect on RSA (Figure 5C).

Our analysis further identified several SNPs located on annotated genes with functions and phenotypes directly linked to RSA. For instance, S7_2952840 (*p* = 6.96E-08) was mapped to a gene annotated as *ILR3-like*. *ILR3* (also known as *bHLH105*) encodes a basic helix-loop-helix (bHLH) transcription factor involved in iron homeostasis (Tissot et al. 2019) putatively though auxin signaling (Rampey et al. 2006). Likewise, S10_11497177 (*p* = 3.08E-08) was found upstream of the *bHLH18*; is associated with stress response and hormone signaling pathways, including ethylene and gibberellins (Cui et al. 2018). Additionally, SNP 54899232 (*p* = 8.77E-09) was positioned in the exon of a gene annotated as a transcription corepressor of *LEUNIG*. Prior studies have shown that the *LEUNIG_HOMOLOG* (*LUH*) plays a critical role in root elongation (Sitaraman et al. 2008).

## Discussion

### Sorghum exhibit variation that adapts to nitrogen and phosphorus deficiency

Root system architecture traits varied significantly among the genotypes. A HC dendrogram based on all RSA analyzed grouped the genotypes into five clusters. However, when N and P were removed, only two clusters emerged. This reduction in the number of clusters could be indicative of differences in root architecture driven by N and P deprivation as previously described (Parra-Londono et al. 2018) whereby RSA types were described as: small; compact and; exploratory root system, all linked to adaptation under N, P or water stress.

### Sorghum adapts to Nitrogen and Phosphorus deficiency using a root elongation-thickness trade-off strategy

PCA, pairwise correlation and Gaussian graphical models and fixed-effects model analyses all showed RSA responded in a nutrient-specific manner.

In PCA analysis, increase in the contribution of PC1 from 56.9% under optimal conditions to 62.6% under N deficiency implies that RSA contributing to the variance under N deficiency (mainly root elongation and number of branch points) increased. Conversely, decrease in the variance in PC2 to 28.0% indicates that the RSA in PC2 (diameter traits) decreased (root elongation-thickness trade-off). This trade-off was more pronounced under P deficiency as displayed by the perpendicular orientation to PC1 to diameter traits in PC2.

Correlation analysis and Gaussian graphical models further highlighting RSA reorganization in a nutrient deficient manner with N removal showing a coordinated modular organization of RSA with the total root length as the hub with negative correlations to diameter measurements. P removal led to a similar switch in RSA but with root length diameter range 2 becoming the central trait and a more pronounced root elongation-thickness trade off – with root elongation being driven by fine root classes (root length diameter ranges 1 and 2).

Fixed-effects model analyses affirmed root elongation-thickness trade-off whereby surface area and volume, declined under both N and P deficiency, with the strongest reductions observed under P limitation.

A similar trade-off was reported described by York et al., (2013). The authors argue that thicker roots (higher root diameter) tend to be less branched or have lower branching frequency, particularly in resource-constrained environments. In our study, we propose that under P deficiency, sorghum reallocates growth toward elongation and proliferation of fine roots, thereby increasing total root length and enhancing soil exploration capacity. Fine roots are metabolically less costly to produce and provide greater absorptive surface per unit biomass, making them advantageous for acquiring immobile nutrients such as phosphorus.

Fine root proliferation under P deficiency is consistent with results obtained in rice (Wissuwa 2003; Shimizu et al. 2004; Yi et al. 2005) and in maize (Mollier and Pellerin 1999); but in contrast to shorter root growth observed in Arabidopsis grown in low P conditions. The varying effects of P deficiency on root development may be attributed to species and genotypic variations, as Reviewed in (Niu et al. 2013). Reduced root secondary growth is consistent with the notion that reduced root secondary growth decreases maintenance and construction costs, allowing greater root elongation and soil exploration, thereby improving P acquisition and plant growth under nutrient stress. The hypothesis was advanced by (Strock et al. 2017). Further support comes from studies of sorghum RSA (Parra-Londono et al. 2018), where increased elongation of lateral roots under P deficiency produced a more highly branched root system.

### GWAS reveal genetic factors associated with root system adaptation under nitrogen and phosphorus deficiency

Our GWAS analysis approach leveraging BLINK, FarmCPU and MLMM approaches proved powerful in identifying SNPs associated with RSA adaptation under N and P deficiency. We chose these approaches because compared to traditional single-locus mixed linear models (MLM), these multi-locus models offer greater statistical power and improved resolution for dissecting complex traits, increasing the likelihood of identifying associations that might otherwise go undetected. FarmCPU achieves this by iteratively separating marker testing from kinship correction, treating significant markers as fixed effects while modeling random background variation, thus reducing false positives (Liu et al. 2016). And, MLMM extends MLM by incorporating significant markers stepwise as cofactors, allowing simultaneous modeling of multiple loci and capturing more complex genetic architectures (Segura et al. 2012).

Expectedly, SNPs were detected based on the surface area, volume, root length diameter range 2 and to a lesser extent branching frequency RSAs. GWAS results revealed a significant SNP that colocalized with surface area and root length diameter range 2. Further analysis of phenotypic variation linked to the different haplotypes of S1_26205071 (C allele vs. T allele) demonstrated the phenotypic impact of this SNP where C Allele accessions exhibited significantly higher values of surface area and root length diameter range 2 . The haplotype analysis confirms that this SNP is not just statistically significant but biologically relevant in shaping important root traits, making it a strong candidate for further validation and incorporation into breeding strategies aimed at improving root system architecture. Although no gene was associated with the SNP, this SNP could be linked to a causative variant in the vicinity, functioning through linkage disequilibrium. Additional fine-mapping or functional studies, such as transcriptome analysis, could reveal specific genes affected by this SNP.

Our GWAS analysis revealed significant associations with additional SNPs located in gene-coding regions, with three noteworthy genes identified: (i) The *ILR3-like* gene belongs to the basic helix-loop-helix (*bHLH*) family of transcription factors, which are involved in iron homeostasis, nutrient acquisition, and root development. In *Arabidopsis thaliana*, the *ILR3* gene regulates iron deficiency responses in roots, which can indirectly influence root growth traits under iron-limiting conditions (Tissot et al. 2019). Given its involvement in iron regulation, *ILR3*-*like* may mediate crosstalk between nitrogen signaling and other nutrients, such as iron. Under conditions of nutrient imbalance, such as nitrogen deficiency, *ILR3*-*like* might trigger structural changes in roots, enhancing branching and improving nutrient foraging efficiency. (ii) *bHLH18*, another member of the bHLH transcription factor family, is associated with diverse plant growth and developmental processes (Cui et al. 2018), including root development, stress responses, and hormonal signaling pathways (e.g., auxin and gibberellin). The association between *bHLH18* and root volume suggests that this gene may influence root growth by regulating cell proliferation, expansion, or overall biomass accumulation in roots. (iii) Transcriptional Repressor of *LEUNIG* (*LUH*) likely acts as a positive regulator of root elongation, influencing cell division and expansion processes in the root meristem or elongation zone (Sitaraman et al. 2008). This hypothesis is further supported by the identification of a SNP linked to LUH using root length diameter range 2 (RLDR2) as a trait, suggesting its role in promoting elongation under specific root system adaptations.

The associations of these genes with RSA traits suggest they may function in the same or complementary pathways that modulate root structure and function. Further functional genetics validation will provide critical insights into their roles and interactions, offering new avenues for understanding and improving root system adaptations to nutrient stress.

In conclusion, our study underscores the vast genetic diversity within the sorghum gene pool, making it a valuable resource for identifying alleles for germplasm improvement. Integrating high-throughput root architecture phenotyping with freely accessible genomic data enabled us to pinpoint genetic regions associated with RSA adaptation under nutrient-limiting conditions and to infer, plausibly the biological processes involved. This knowledge may inform crop breeding strategies designed to improve resilience to challenges associated with soil nutrient deficiency.

**Table 1.** Sorghum single nucleotide polymorphisms (SNPs) showing significant genome-wide associations with root system architecture (RSA) traits (FDR correction at ≤0.05). SNPs within genes are presented with their annotations in order of chromosomal location. AF: allele frequency; AdjP: P-value after FDR adjustment. *Represent SNPs identified under nitrogen deficiency, ^†^represent SNPs identified under phosphorus deficiency. Letters after SNP positions represent GWAS model used: a = BLINK, b = FarmCPU, c = MLMM.

| Chr | Position | AF | AdjP | Associated Trait | Gene ID | Annotation |
| --- | --- | --- | --- | --- | --- | --- |
| 1 | 26205071<sup>*c</sup> | 0.16 | 6.18E-08 | Surface area | N/A | N/A |
| 1 | 26205071<sup>*c</sup> | 0.16 | 1.67E-07 | Volume | N/A | N/A |
| 1 | 26205071<sup>*b</sup> | 0.16 | 4.05E-08 | Surface area | N/A | N/A |
| 1 | 26205071<sup>†c</sup> | 0.16 | 1.89E-13 | RLDL2 | N/A | N/A |
| 1 | 26205071<sup>*c</sup> | 0.16 | 5.43E-08 | Surface area | N/A | N/A |
| 1 | 26205071<sup>*c</sup> | 0.16 | 5.43E-08 | Surface area | N/A | N/A |
| 1 | 26205071<sup>*c</sup> | 0.16 | 1.42E-09 | RLDL2 | N/A | N/A |
| 1 | 26423820<sup>*c</sup> | 0.20 | 1.05E-07 | Volume | N/A | N/A |
| 1 | 26731398<sup>*c</sup> | 0.13 | 1.78E-07 | Surface area | N/A | N/A |
| 1 | 26731398<sup>*c</sup> | 0.13 | 1.72E-08 | Volume | N/A | N/A |
| 1 | 26731398<sup>*b</sup> | 0.13 | 1.19E-07 | Surface area | N/A | N/A |
| 1 | 26731398<sup>*b</sup> | 0.13 | 3.70E-11 | Volume | N/A | N/A |
| 1 | 26731398<sup>*c</sup> | 0.13 | 3.29E-08 | Volume | N/A | N/A |
| 1 | 27129528<sup>*c</sup> | 0.10 | 2.20E-08 | Volume | XM_002464586.2 | CCRP 4 homolog |
| 1 | 26205071<sup>*a</sup> | 0.16 | 1.10E-10 | Surface area | N/A | N/A |
| 1 | 26423820<sup>*a</sup> | 0.20 | 4.68E-08 | Volume | N/A | N/A |
| 1 | 26205071<sup>*a</sup> | 0.16 | 1.56E-12 | RLDL2 | N/A | N/A |
| 1 | 20102651<sup>*b</sup> | 0.40 | 8.66E-11 | Volume | N/A | N/A |
| 2 | 63695738<sup>†c</sup> | 0.42 | 2.03E-08 | RLDL2 | N/A | N/A |
| 2 | 63695779<sup>†b</sup> | 0.41 | 5.28E-09 | RLDL2 | N/A | N/A |
| 3 | 15123886<sup>*b</sup> | 0.10 | 1.97E-08 | RLDL2 | XP_002455526 | Uncharacterized protein |
| 3 | 54899232<sup>†c</sup> | 0.08 | 8.77E-09 | RLDL2 | XP_02131268.1 | LEUNIG |
| 3 | 67879725<sup>*a</sup> | 0.12 | 6.48E-11 | Volume | XP_002456657.1 | Cell Division Control Protein 6 |
| **4** | 12760823<sup>†b</sup> | 0.28 | 6.82E-11 | RLDL2 | N/A | N/A |
|  | 5702394<sup>*b</sup> | 0.09 | 8.74E-08 | NBP | XP_002451690.2 | Uncharacterized protein |
| **4** | 58775820<sup>*b</sup> | 0.16 | 3.12E-09 | NBP | XP_021314639.1 | *MAPK14-like* |
| **5** | 17754110<sup>*b</sup> | 0.04 | 3.38E-08 | NBP | XP_002449381.2 | 1-aminocyclopropane-1-carboxylate oxidase 1 |
| **5** | 59062428<sup>*b</sup> | 0.12 | 2.63E-08 | Volume | N/A | N/A |
| **6** | 55153514<sup>†b</sup> | 0.08 | 6.48E-08 | RLDL2 | XP_021318054.1 | E3 ubiquitin-protein ligase RNF14 |
| **6** | 56342501<sup>*c</sup> | 0.03 | 2.55E-08 | N NBP | XP_021318339.1 | probable LRR receptor-like serine/threonine-protein kinase/hypothetical protein |
| **6** | 56929524<sup>*b</sup> | 0.03 | 4.41E-09 | Volume | SORBI_3006G223101 | Hypothetical protein |
| **6** | 4077580<sup>*a</sup> | 0.05 | 1.16E-09 | NBP | XP_021318130.1 | Uncharacterized protein |
| **6** | 45806791<sup>*b</sup> | 0.27 | 1.29E-10 | Volume | XP_002446521.1 | Transcription factor bHLH18 |
| **7** | 2952840<sup>*c</sup> | 0.07 | 6.96E-08 | NBP | XP_0213120712.1 | ILR3 |
| **7** | 64170934<sup>*b</sup> | 0.43 | 3.75E-13 | Volume | N/A | N/A |
| **8** | 15675606<sup>*b</sup> | 0.05 | 3.96E-09 | Volume | N/A | N/A |
| **9** | 49587358<sup>†b</sup> | 0.35 | 1.35E-11 | NBP | N/A | N/A |
| **9** | 49587358<sup>*a</sup> | 0.35 | 1.44E-09 | NBP | N/A | N/A |
| **10** | 9879684<sup>*b</sup> | 0.16 | 5.00E-09 | NBP | XP_002438177.2 | peroxygenase 4 |
| **10** | 17059423<sup>*a</sup> | 0.35 | 2.99E-09 | Volume | N/A | N/A |
| **10** | 11497177<sup>*b</sup> | 0.46 | 3.08E-08 | Volume | N/A | N/A |

## Author contributions

SR and second author JM conceived and designed the study. First author JM performed the experiments and all analysis guided by SR and CM. SR, and first author JM, wrote the manuscript. All authors read and approved the final manuscript.

## Data availability statement

The data that support the findings were derived from the following resources available in the public domain: [https://www.morrislab.org/data and https://genebank.icrisat.org/Common/Viewer?ctg=Referenceset&ref=RS1]. The data that supports the findings of this study are available in the supplementary material of this article. Dataset S1 contain the sorghum accessions metadata and Dataset S2 contain the root system architecture (RSA). Data of sorghum accessions used in the study. Full results from the GWAS, heritability estimates, PCA analysis, and all associated scripts used in the study have been uploaded to the Genetics Society of America’s Figshare portal.

## Acknowledgements

This work was funded by the National Research Fund grant to S.R. contract number KU/DVCR/NRF/VOL1/27 under the Multi Disciplinary Research Grants. The authors thank and acknowledge the International Crops Research Institute for the Semi-Arid Tropics (ICRISAT) through Dr. Eric Manyasa for availing the sorghum germplasm used in this project.

## Supplemental information

The following Supporting Information is available for this article:

**Table S1**: Description of root system architecture (RSA) traits extracted from RhizoVision analysis.

**Table S2.** Genetic and environmental effects on sorghum root system architecture.

**Figure S1.** Genetic diversity of sorghum accessions used for sorghum root system architecture analysis.

**Figure S2.** Sorghum root clusters generated under different nitrogen and phosphorus deficiency.

**Figure S3.** Pairwise scatterplot matrix of sorghum root system architecture.

**Figure S4:** Genome-wide marker distribution, quality assessment, and linkage disequilibrium decay

**Figure S5.** Multivariate detection of Single Nucleotide Polymorphisms (SNPs) associated with Root System Architecture (RSA) using multiple traits and models

**Dataset S1.** Sorghum accessions metadata.

**Dataset S2.** Root system architecture data of sorghum accessions used in the study.

**Dataset S3.** PCA loading and variance contributions of sorghum root system architecture

**Dataset S4.** Table of fixed effects and model summary of sorghum root system architecture

