## Supplementary Tables and Figures for "Genomics of Root System Architecture Adaptation in Sorghum under Nitrogen and Phosphorus Deficiency"

The following Supporting Information is available for this article:

**Table S1**: Description of root system architecture (RSA) traits extracted from RhizoVision analysis.

**Table S2.** Genetic and environmental effects on sorghum root system architecture.

**Figure S1.** Genetic diversity of sorghum accessions used for sorghum root system architecture analysis.

**Figure S2.** Sorghum root clusters generated under different nitrogen and phosphorus deficiency.

**Figure S3.** Pairwise scatterplot matrix of sorghum root system architecture.

**Figure S4:** Genome-wide marker distribution, quality assessment, and linkage disequilibrium decay

**Figure S5.** Multivariate detection of Single Nucleotide Polymorphisms (SNPs) associated with Root System Architecture (RSA) using multiple traits and models

**Dataset S1.** Sorghum accessions metadata.

**Dataset S2.** Root system architecture data of sorghum accessions used in the study.

**Dataset S3.** PCA loading and variance contributions of sorghum root system architecture

**Dataset S4.** Table of fixed effects and model summary of sorghum root system architecture

**Figure S1.** Genetic diversity of sorghum accessions used for root system architecture analysis. (A) Map showing the collection locations of the sorghum accessions. (B) A neighbor-joining (NJ) tree illustrating the genetic structure and relationships among sorghum races. (C) An admixture plot depicting the clustering patterns of sorghum races at the optimal K value (K = 7). The key shows color codes for sorghum races on the map and NJ tree. The key provides color codes for sorghum races shown on the map and the NJ tree.

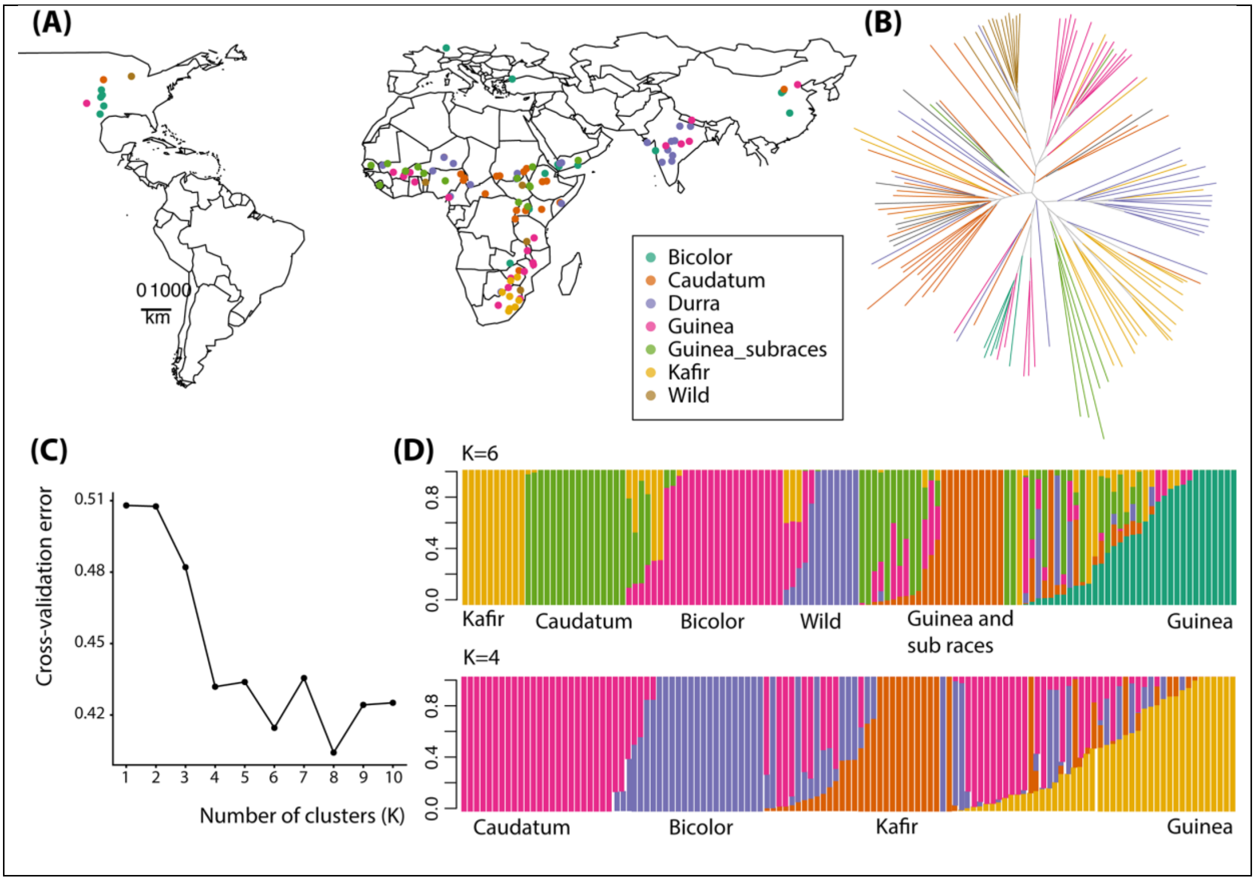

**Table S1**: Description of root system architecture (RSA) traits extracted from RhizoVision analysis (Seethepalli et al., 2020).

| Trait | Abbreviation | Category | Description |
| --- | --- | --- | --- |
| Total root length | TRL | Size | Computed by determining the Euclidean distance of the medial axis pixel in the skeletonized image. |
| Perimeter | P | Size | Total Euclidean distance of contour pixels in the segmented image |
| Average diameter | AD | Distribution | For each pixel in the skeletonized image, the distance to the nearest non-root pixel is computed. This distance is used as radius to fit a circle. The diameter of the circle at each pixel is noted as the root diameter. The list of diameter from all the medial axis is used to determine the average median and maximum diameter for the entire root crown. |
| median diameter | MD | Distribution | |
| Maximum diameter | MaxD | Distribution | |
| Branch points | NBP | Extent | Computed by counting the total number of branch pixels in the skeletonized image with topology. |
| Branching frequency | BF | Extent | The number of branch points divided by the total root length. |
| Number of Root Tips | NRTP | Extent | Count of total number of tip pixels in the skeletonized image. |
| Volume | V | Size | Using the radii determined earlier, the sum of all cross-sectional areas across all the medial axis pixels is noted as the volume and the sum of the perimeter across all the medial axis pixels is noted as the surface area. |
| Surface area | SA | Size | |
| Network Area | NA | Size | Total number of pixels in the segmented image. |
| Root length Diameter range 1 | RLDR1 | Distribution | Fine roots below 2 mm or separate lateral roots from their parents through interactive adjustment. |
| Root length Diameter range 2 | RLDR1 | Distribution | Separate lateral roots from RLDR1 through interactive adjustments. |

**Table S2. Genetic and environmental effects on root system architecture**. The interaction between genotypes and the environment was significantly different for maximum diameter (MaxD), number of branching points (NBP), branching frequency (BF), number of root tips (NRTP), volume (V), root length diameter range A (RLDRA), root length diameter range B (RLDRB) (Table 4.1). Heritability was highest in all root traits, with lower heritability observed in maximum diameter and branching frequency. ANOVA results for genotypes (Gb) at different nutrient conditions and environment, and genotype × environment interactions (G × E) at deprived nitrogen and deprived phosphorus. * $p < .05$. **$p < .01$, ***$p < .001$: ns not significant.

| **Trait** | | **All** | | | **N-Dep** | | | **P-Dep** | | | **Anovab** | | **$h^2$** |
|---|---|---|---|---|---|---|---|---|---|---|---|---|---|
| | Mean | s.d. | Gb | Mean | s.d. | Gb | Mean | s.d. | Gb | E | G | G×E | |
| **TRL** | 2110.69 | 1117 | *** | 1996.54 | 1297.47 | *** | 2025 | 1154.83 | *** | *** | *** | ns | 0.9479 |
| **P** | 3215.17 | 1502.8 | *** | 3136.71 | 1710.99 | *** | 3180.5 | 1627.03 | *** | * | *** | ns | 0.9491 |
| **AD** | 1.87 | 0.64 | *** | 1.68 | 0.52 | *** | 1.76 | 0.6 | *** | *** | *** | ns | 0.9389 |
| **MD** | 1.66 | 0.57 | *** | 1.5 | 0.47 | *** | 1.58 | 0.54 | *** | *** | *** | ns | 0.9396 |
| **MaxD** | 7.29 | 2.02 | *** | 6.95 | 1.5 | *** | 6.89 | 1.65 | *** | *** | *** | *** | 0.8725 |
| **NBP** | 381.95 | 258.89 | *** | 360.51 | 360.67 | *** | 357.28 | 278.18 | *** | *** | *** | *** | 0.9128 |
| **BF** | 0.16 | 0.05 | *** | 0.15 | 0.05 | *** | 0.16 | 0.05 | *** | ns | *** | *** | 0.8594 |
| **NRTP** | 50.4 | 19.64 | *** | 50.18 | 20.7 | *** | 52.77 | 21.94 | *** | ns | *** | *** | 0.9095 |
| **V** | 8079.75 | 6608.85 | *** | 5752.31 | 4879.87 | *** | 6365.69 | 4903.07 | *** | *** | *** | *** | 0.8976 |
| **SA** | 12163.11 | 7622.28 | *** | 10120.24 | 7105.57 | *** | 10679.73 | 6572.01 | *** | ** | *** | ns | 0.9445 |
| **NA** | 2921.66 | 1691.94 | *** | 2500.37 | 1543.19 | *** | 2636.17 | 1494.9 | *** | ns | *** | ns | 0.95 |
| **RLDR1** | 1057.44 | 644.46 | *** | 1119.53 | 795.84 | *** | 1100.26 | 740.34 | *** | *** | *** | *** | 0.904 |
| **RLDR1** | 1053.25 | 604.09 | *** | 877.01 | 605.9 | *** | 924.74 | 558.35 | *** | ns | *** | *** | 0.9063 |

**Figure S2.** Sorghum root clusters generated under different nutrient conditions. The average silhouette width is plotted against the number of clusters (k) to determine the optimal number of clusters for each condition: Top (Optimum): Clustering under optimal conditions shows the highest average silhouette width at k = 5, indicating five distinct RSA clusters. Middle (N-Dep): Clustering under nitrogen deficiency (N-Dep) shows the highest silhouette width at k = 2, suggesting two major RSA clusters. Bottom (P-Dep): Clustering under phosphorus deficiency (P-Dep) indicates an optimal cluster number of k = 2 based on the highest silhouette width.

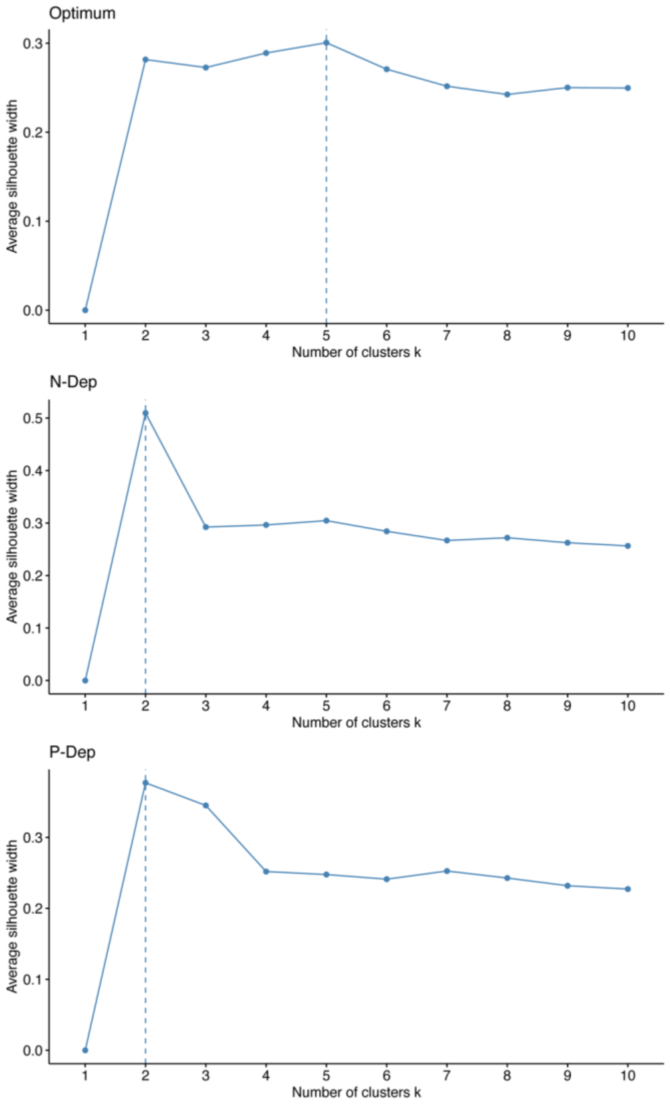

**Figure S3.** Pairwise scatterplot matrix of root system architecture traits. Diagonal panels depict trait distributions. Lower panels show scatterplots of raw trait values, and upper panels present Pearson correlation coefficients ($r$) with associated significance levels indicated by asterisks ($P < 0.05$, P < 0.01, $P < 0.001$).

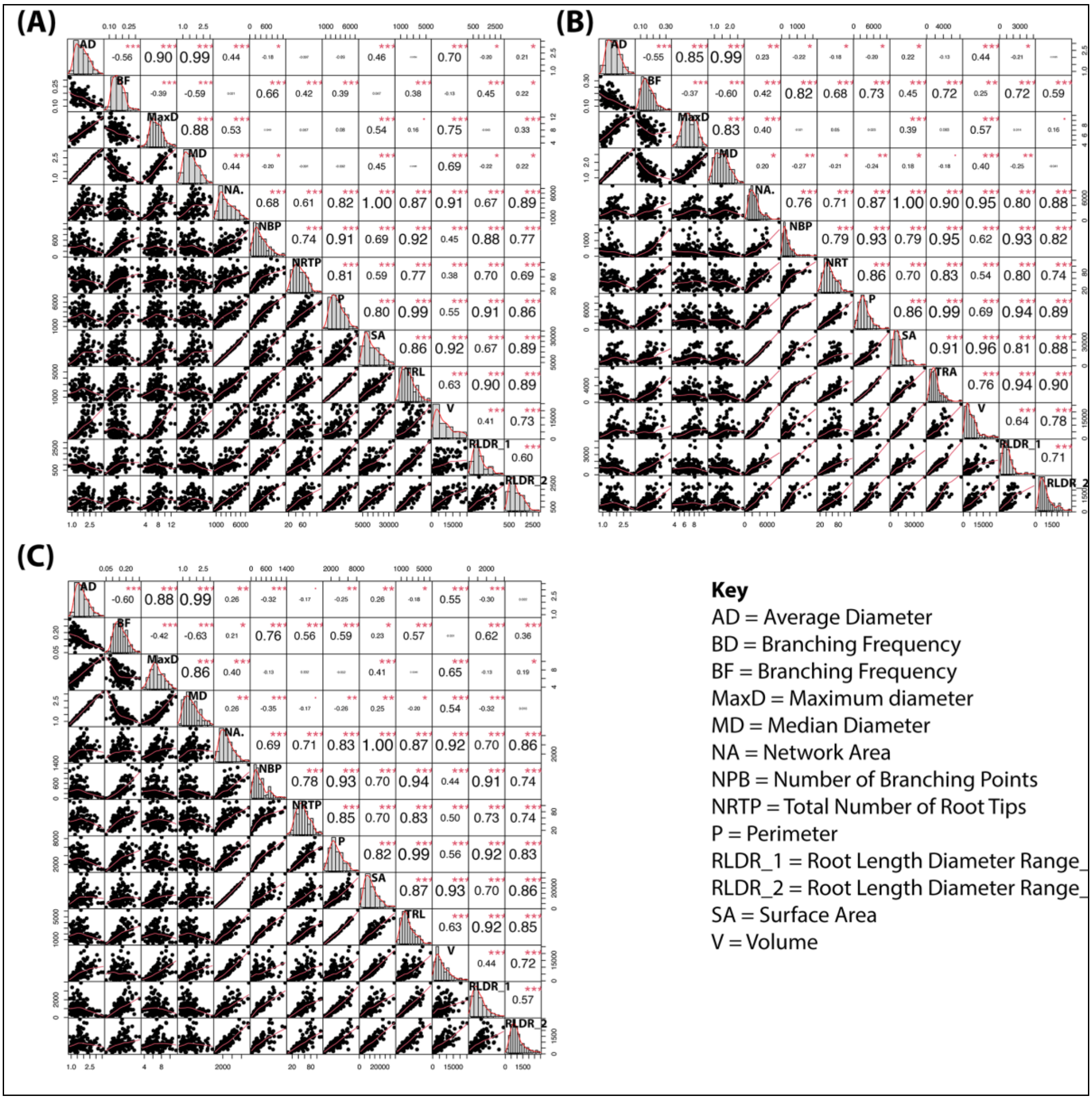

**Figure S4:** Genome-wide marker distribution, quality assessment, and linkage disequilibrium decay. (A) Single nucleatide polymorphism (SNP) density across the genome calculated in 50 kb windows. Color intensity represents the number of SNPs per 0.05 Mb window, illustrating overall marker coverage across chromosomes 1–10. (B) Linkage disequilibrium decay estimated as pairwise $r^2$ plotted against physical distance (kb) for each chromosome. LD declines rapidly within the first 100 kb and gradually approaches background levels at larger distances. (C) Marker quality across the genome summarized as mean missing genotype rate per 50 kb window. Most genomic regions exhibit low to moderate missingness, indicating good data quality. (D) Minor allele frequency (MAF) distribution across the genome, shown as mean MAF per 50 kb window.

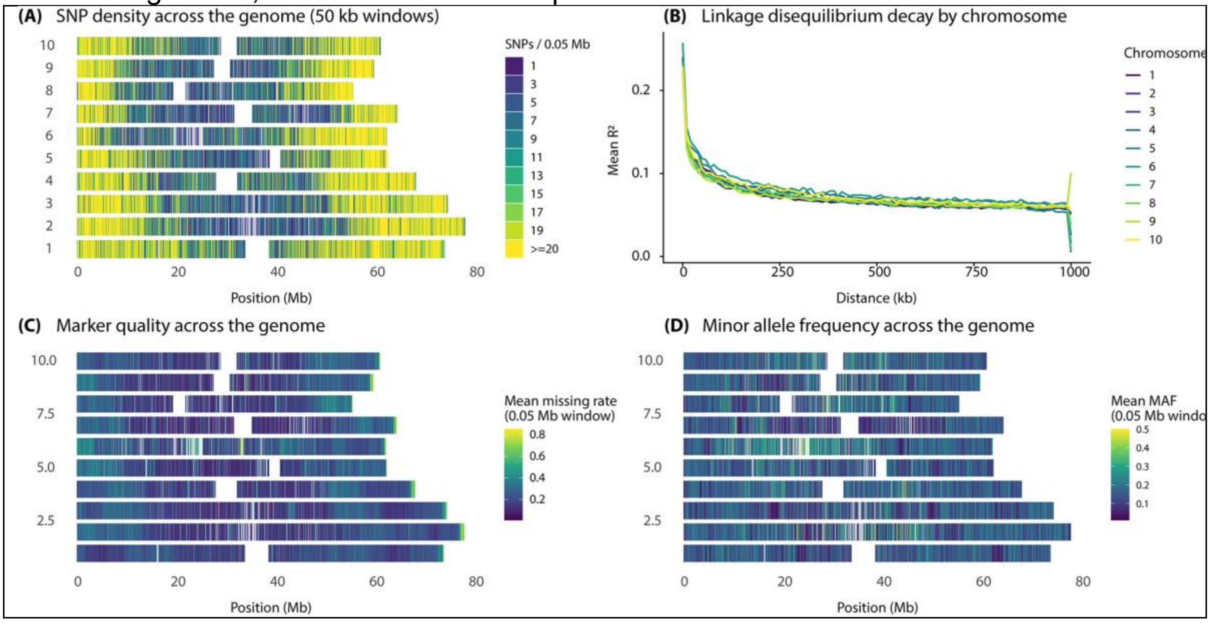

**Figure S5.** Multivariate detection of Single Nucleotide Polymorphisms (SNPs) associated with Root System Architecture (RSA) using multiple traits and models. **A**. Manhattan Plots: The plots highlight the most prominent SNP, S1_26205071, identified using the Fixed and Random Model Circulating Probability Unification model based on four RSA traits: root length diameter range 1 (RLDR1), volume, surface area, and number of branching points. Additionally, SNP S1_26731398 was detected in both the Multi-Locus Mixed Model and Bayesian-information and Linkage-disequilibrium Iteratively Nested Keyway models using the RLDR1 trait. **B**. Quantile-Quantile (Q-Q) Plots: **Bi**: Q-Q plot for the multivariate analysis performed using the FarmCPU model. **Bii**: Combined Q-Q plot for results obtained from the all models used.

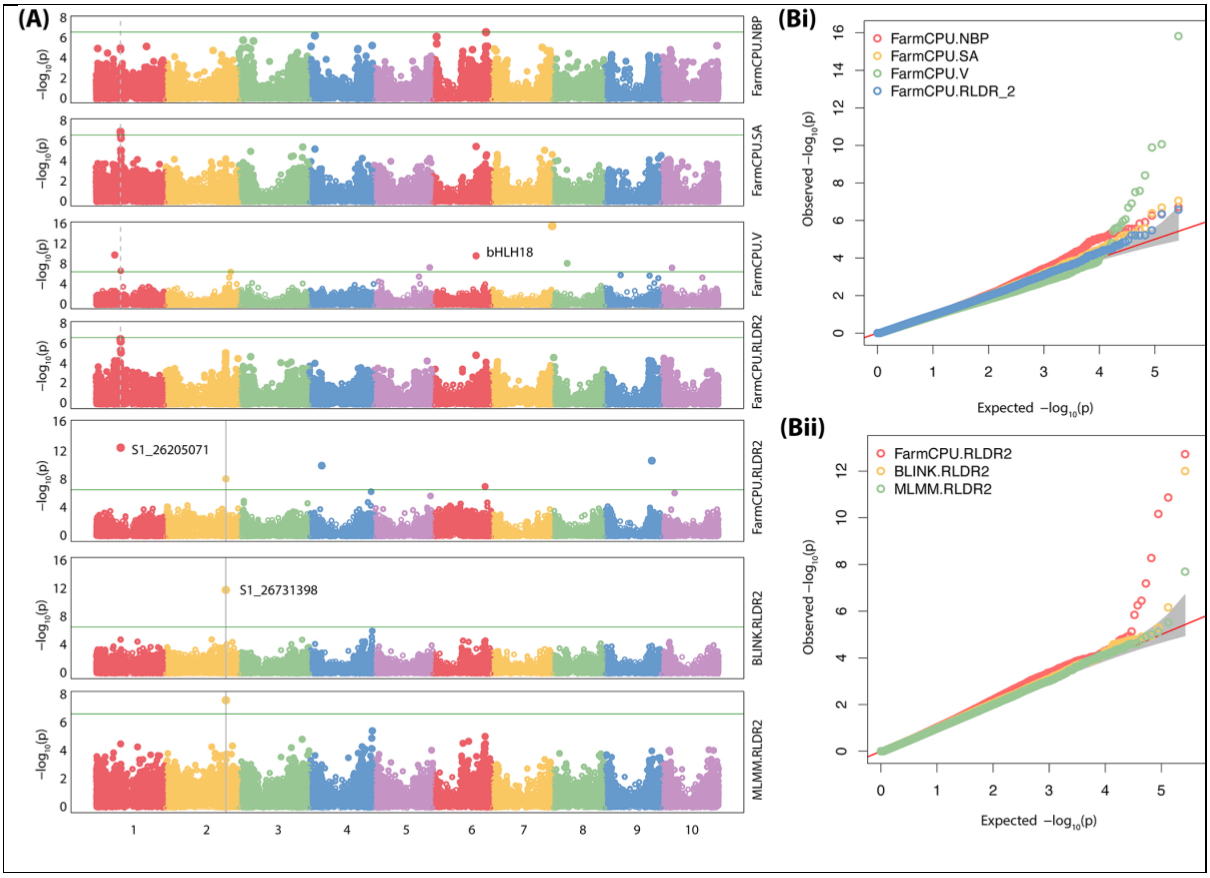
